# Cytokinin Dehydrogenase in Xylem Sap Reveals A Direct Link Between Cytokinin Metabolism and Long-Distance Transport

**DOI:** 10.1101/2023.11.06.565614

**Authors:** Daniel Nedvěd, Václav Motyka, Ivana Raimanová, Petre I. Dobrev, David Kopečný, Pierre Briozzo, Federica Brunoni, Karel Müller, Klára Hoyerová

## Abstract

Metabolic degradation of plant hormones cytokinins (CKs) co-regulates their homeostasis and signalling. In this work, we employed a large-scale bioinformatical analysis to address a diversity of cytokinin oxidase/dehydrogenase (CKX) substrate specificities previously described in several case studies. We present a three-way correlation of the entire CKX amino acid sequences, a variable motif involved in substrate binding, and subcellular localizations predicted by a deep learning model. This correlation is conserved in monocotyledonous plants, suggesting that the CKX diversity in a single species allows a precise tuning of the CK homeostasis. Following these findings, we detected CKX activity in xylem sap for the first time, using the oat (*Avena sativa*) as a model plant. Further investigation of the substrate specificity and glycosylation of this xylem-located CKX suggested that it originates in roots. We have identified 27 putative CKXs in oats and attributed the xylem-located activity to the extracellular isoforms AsCKX1a,c,d. Finally, we show that the xylem-located CKX activity responds to the nitrate supply, highlighting its physiological relevance. Taken together, we show that CKX directly modulates root-to-shoot CK translocation through metabolic degradation of the transported CKs.

## Introduction

Cytokinins (CKs) are a group of structurally related molecules derived from adenine through substitution on the N6 atom. They act as plant hormones – a subset of plant signalling molecules regulating diverse aspects of plant growth and development (Kieber and Schaller, 2014; Wybouw and De Rybel, 2019; Hu and Shani, 2023). Among others, CKs are involved in the complex system mediating the acquisition and distribution of nitrate, a nitrogen source for plants (Abualia, Riegler and Benkova, 2023). Long-distance transport of CKs from roots to shoots via xylem bears information about nitrate availability in soil, and CKs transported this way are responsible for nitrate assimilation in the shoot (Kiba *et al*., 2011; Poitout *et al*., 2018; Roy, 2018; Sakakibara, 2021). CKs further modulate root architecture and expression of nitrate transporters, thus affecting nitrate uptake efficiency (Kiba *et al*., 2011; Kiba and Krapp, 2016; Varala *et al*., 2018; Jia and Von Wirén, 2020; Tessi *et al*., 2020, 2023). Simultaneously, plants regulate CK distribution and activities of enzymes involved in CK biochemistry in response to nitrate availability (Takei *et al*., 2001, 2004; Maeda *et al*., 2018; Poitout *et al*., 2018).

The proper CK signalling depends on processes both maintaining and altering CK homeostasis – transport and conversions between biologically active (i.e. capable of binding to their receptors) and inactive CK forms (reviewed by Hluska et al., 2021; Hu and Shani, 2023; Kieber and Schaller, 2014; Nedvěd et al., 2021). The latter consists of a series of biochemical reactions, of which this paper focuses on CK oxidative degradation catalyzed by cytokinin dehydrogenase (CKX; EC 1.5.99.12; also known as cytokinin oxidase or cytokinin oxidase/dehydrogenase). The CKX reaction involves two steps. Firstly, the CK forms a stable oxidized intermediate with an extra double bond compared to the reactant (Popelková *et al*., 2006; Kopečný *et al*., 2008, 2016). An electron pair moves from the reactant to the cofactor flavin adenine dinucleotide (FAD) covalently bound to the CKX apoenzyme (Frébortová *et al*., 2004; Malito *et al*., 2004). Secondly, the intermediate undergoes hydrolysis, yielding adenine (or its conjugated form, such as adenosine) and an aldehyde derived from the original N6 substituent (Pačes, Werstiuk and Hall, 1971; Brownlee, Hall and Whitty, 1975; Hare and van Staden, 1994). CKX thus cleaves the bond between the N6 atom and its substituent.

CKX amino acid sequences contain several residues directly involved in cofactor or substrate binding. Their mutations gravely affect the catalytic properties of the enzyme. A histidine residue found in a conserved motif GSH binds the FAD cofactor and is essential for enzyme activity and stability (Kopečný *et al*., 2016). An aspartate residue from a conserved motif WTDYL subtracts a hydride from the substrate in the catalytic process (Malito *et al*., 2004; Kopečný *et al*., 2016).

Several other residues co-determine the shape of the enzyme’s binding cavity and provide differential accommodation for various CK species (Kopečný *et al*., 2016). A semi-conserved motif FL**X**RV**XXX**E (with “**X**” denoting a variable residue) is found along the entrance to the binding cavity, and its penultimate residue accounts for the substrate specificities of different CKX enzymes. A glutamate residue in this position interacts with N9 atoms of CK nucleobases and stabilizes them in the binding cavity. Glutamate-containing variants of CKX, such as maize CKX1 (ZmCKX1) or mouse-ear cress CKX2,4 (AtCKX2,4), thus strongly prefer CK nucleobases and also display relatively high enzyme activities *in vitro.* Conversely, when the non-conserved residue is less bulky or polar, the corresponding CKXs are more promiscuous towards CK ribosides and monophosphates, and their enzyme activities are lower (Malito *et al*., 2004; Frébortová *et al*., 2007; Galuszka *et al*., 2007; Kopečný *et al*., 2016). Other authors have also shown that CKX substrate specificities and enzyme activities correlate with the respective enzyme’s subcellular localizations (Šmehilová *et al*., 2009; Kowalska *et al*., 2010; Zalabák *et al*., 2014). Considering sequences of the differentially localized CKXs isoforms studied in these works, one can see correlations among the subcellular localizations and the identity of the variable ligand-binding residue mentioned above.

CKX enzymes also participate in the cross talk of CKs and nitrogen (Reid *et al*., 2016; Gigli-Bisceglia *et al*., 2018). Given the specific roles of different CKX forms reported in the works cited earlier, it has intrigued us whether it is possible to pinpoint CKXs regulating the amount and quality of CKs transported from roots to shoots in response to the nitrogen availability in the environment. CKX-mediated regulation of CK long-distance flux has already been suggested (Yang, Yu and Goh, 2002; Brugière *et al*., 2003; Hoyerová *et al*., 2007), but the issue has remained unresolved until nowadays. Based on preliminary data, we hypothesize that some CKX proteins are active in the xylem sap and directly affect CKs transported from roots to shoots (Hoyerová *et al*., 2007).

Connecting this CKX activity to a CKX isoform with particular substrate specificity would allow us to discuss how CK degradation shapes their long-distance flux (and thus information about the nitrogen availability).

## Results

### CKX Diversity Reflects in the Amino Acid Composition of A Non-Conserved “Entrance” Triplet

Before exploring the relationship between CKX activity and the long-distance transport of CKs, we performed a large-scale bioinformatic analysis of CKX sequences to establish their classification according to their biochemical properties. Of the whole plant kingdom, we focused on monocotyledons, which had proven to be good model plants in our previous studies concerning CK degradation in xylem sap (Hoyerová *et al*., 2007).

To determine whether sequentially similar CKXs share common identities of the ligand-binding variable motifs, we considered 492 CKX sequences from various monocotyledonous species. For their complete list, see Table S1. Through multi-sequential alignment and subsequent dendrogram construction, we identified eight major clusters of sequentially close CKXs (Figure 1). For this work, we named these clusters CKX classes I-VIII. Each class can be characterized by a specific predominant composition of the non-conserved triplet within the FL**X**RV**XXX**E motif (see Introduction). These characteristic triplets are H**X**S (class I), NRV (class II), RME (class III), H**X**G (class IV), **XX**E (class V), HGE (class VI), RDG (class VII), and HKA (class VIII). Notably, the third residue, capable of direct interaction with the CKX’s substrate, is, in most cases, a valine, glutamate, glycine, alanine, or serine. For simplicity, we will refer to the non-conserved triplet as the “entrance triplet” (given its position within the protein) and its third residue as the “VEGAS residue” (based on the five most abundant amino acids).

**Figure 1:**
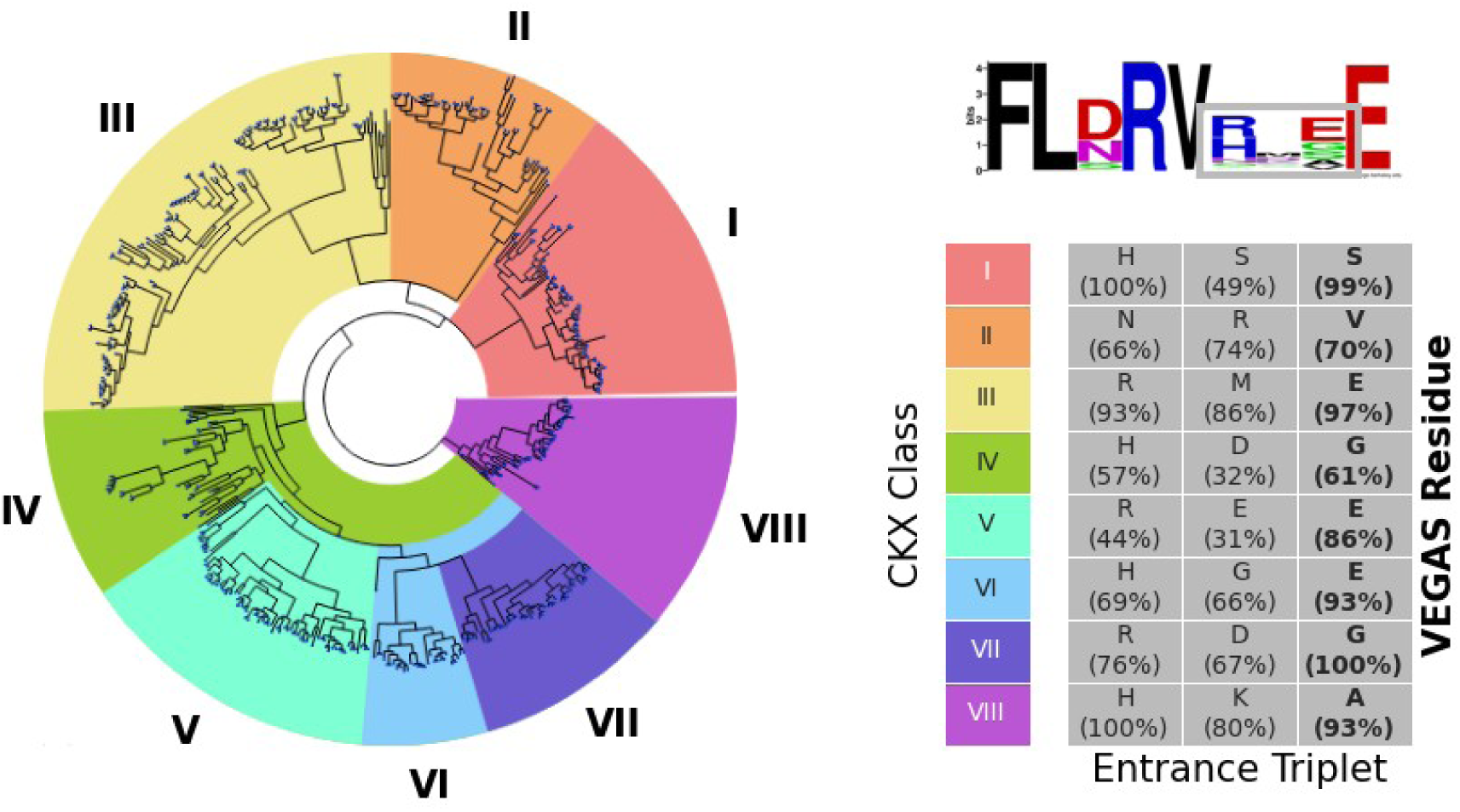
Sequence-based classification of CKXs. **Left**: A tree graph of monocot CKX protein sequences. Each leaf represents a single sequence. The coloured portions of the tree represent eight CKX classes established in this work, annotated as I-VIII. **Top right**: The sequence logo of the semi-conserved FL**X**RV**XXX**E motif with the “entrance triplet” marked by a grey rectangle. Created with WebLogo (Crooks et al., 2004). **Bottom right**: The most common amino acids at each position of the entrance triplet per each CKX class. The occurrence of the residues at the given position and in the given classes are presented in parentheses. If the occurrence of the most common residue is less than 50 %, it is considered variable and labelled “**X**” in the main text. The third position, depicted in the bold font, is the VEGAS residue, which can directly interact with the substrate.

### Crystal Structure of A Class VII Maize CKX Supports the Role of VEGAS Residue in Ligand Recognition

The contribution of VEGAS residue on the shape the CKX active site and its capability to interact with the substrate (or the lack thereof) can be observed on the enzyme’s crystal structures. Up to date, the publicly available structures include those of class I ZmCKX2 (Kopečný *et al*., 2016), class V ZmCKX8 (Nisler *et al*., 2021), class VI ZmCKX1 (Malito *et al*., 2004) and class VIII ZmCKX4a (Kopečný *et al*., 2016). Crystal structures of AtCKX7 (Bae *et al*., 2008) and flax CKX7 (LuCKX7) (Wan *et al*., 2019) are available as well; these two isoforms weren’t part of our analysis, but they show sequential similarity to class II CKXs.

Here we present the structural analysis of class VII ZmCKX5 (refined up to 1.65 Å resolution) comprising glycine residue at the entrance to the active site equivalent to the VEGAS position. The structure is available via Protein Data Bank (PDB) under the code 8QVT. Refinement statistics is given in Table S2. The structure of ZmCKX5 is very similar to those of ZmCKX1 and 8 with root-mean-square deviation (RMSD) values of ∼ 1.0 Å and sequence identity of 59 %.

ZmCKX5 displays a classical two-domain topology and carries a covalently linked FAD (Figure 2). The protein is more ordered at the N-terminus where an additional helix can be observed. The major difference concerns the region of residues 312-350, composed of two helices and a loop in the substrate binding domain. The corresponding regions are well defined in ZmCKX1 and 8 but disordered in the high-resolution structure of ZmCKX5 and all available structures of ZmCKX4a. In ZmCKX2, AtCKX7 and LuCKX7, the region is ordered but adopts several different conformations.

**Figure 2:**
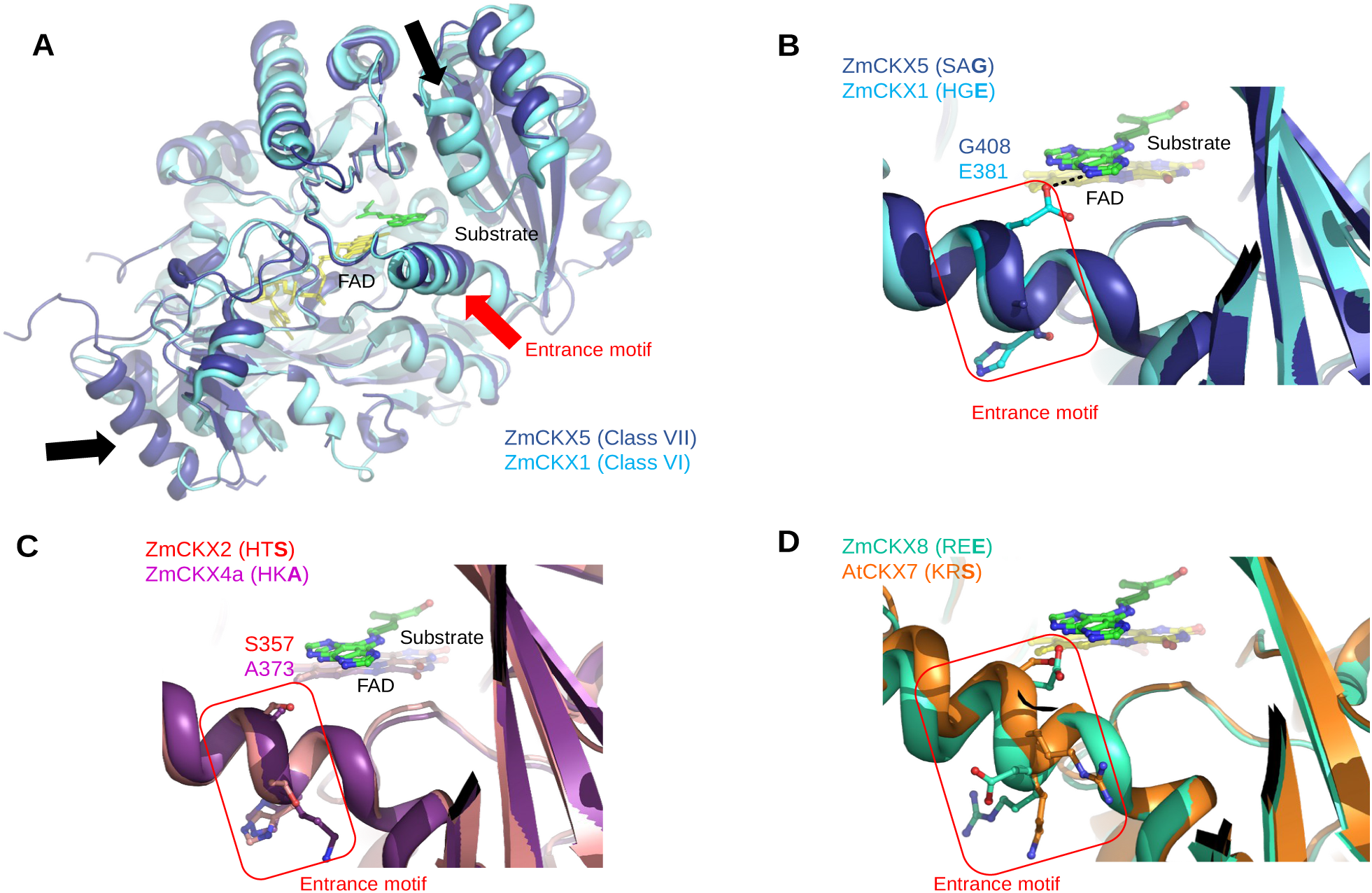
Structure-based substrate specificities of CKXs from different classes. **A**: A structural comparison between ZmCKX5 (dark blue, this work / PDB 8QVT) and ZmCKX1 (light blue, PDB 1W1R). The helix–loop–helix region from residues 294–325 in ZmCKX1 (shown with black arrow) is disordered in ZmCKX5. An additional N-terminal helix in ZmCKX5 is also shown. **B-D**: Detailed overviews at the entrance to the active site in CKXs from six classes. Superpositions of ZmCKX5 on ZmCKX1, ZmCKX2 (red, PDB 4ML8) on ZmCKX4a (violet, PDB 4O95), and ZmCKX8 (cyan, PDB 6YAQ) on AtCKX7 (orange, PDB 2EXR) are shown. The entrance motif is in the red rectangle, and the VEGAS residues are labelled.

Several residues from this region (N303 and T304 in ZmCKX1) are just above the entrance triplet, and together they delineate the substrate entrance comprising residues surrounding the ribosyl or glucosyl moiety of the N9-substitued substrates. It has been previously described that a point mutation at the VEGAS position is not enough to switch the specificity observed among classes, and it has been hypothesized that the variations in the region above this residue have to contribute to the substrate preferences as well (Kopečný *et al*., 2016).

ZmCKX5 crystallized as a covalent homodimer linked through an intermolecular disulphide bond formed by the protomers’ C82 residues. The multi-sequential alignment of the CKX sequences described above revealed that the cysteine residue involved in the disulphide linkage is conserved in approximately 70 % of class V CKXs. By contrast, its occurrence does not exceed 4 % in either of the other classes. Nevertheless, the gel permeation chromatography revealed that ZmCKX5 is a monomeric enzyme in a solution at a neutral pH and an ionic strength of 0.15. A calculation performed by PISA (Krissinel and Henrick, 2007) predicted that only 3.3 % of the solvent-accessible surface area is involved in the dimer interface formation. The gain in solvation free energy upon the interface formation was only –3.5 kcal mol^-1^, and the free energy of assembly dissociation was negative (–1.3 kcal mol^-1^), both indicating instability of the dimer. However, the dimer calculations fall within a “grey region” of complex formation criteria, and the *in vivo* existence of the dimer thus cannot be excluded.

### CKX Classes Clustering Correlates with Predicted Subcellular Localizations

The diversity of CKX substrate specificity has been related not only to the VEGAS residue but also to the subcellular localization of proteins (Šmehilová *et al*., 2009; Kowalska *et al*., 2010; Zalabák *et al*., 2014). In the context of the results above, these findings suggest that different CKX classes, each with their substrate preferences, might also act in specific cellular compartments. So, as the next step, we investigated the correlations between the CKX classification and subcellular localization.

Due to the sparse knowledge of the actual subcellular localizations of the CKX proteins in our dataset, we had them predicted by DeepLoc software (Thumuluri *et al*., 2022). For each sequence, we obtained prediction scores for ten compartments. To visualize correlations between the CKX classification and their predicted subcellular localizations, we transformed the DeepLoc scores from 10-dimensional to 2-dimensional space using the t-distributed stochastic neighbour embedding (tSNE) method (van der Maaten and Hinton, 2008). Figure 3A shows the CKX sequences in the transformed space. The plot reveals that class I (H**X**S), II (NRV), VI (HGE), and VIII (HKA) CKXs form distinct clusters. It means they are likely localized in the same compartment. Notably, the class II cluster lies far from the others in the plot, suggesting unique subcellular localization. Less coherent clusters are formed by class III (RME), V (**XX**E), and VII (RDG) CKXs. Finally, class IV CKXs (H**X**G) do not form any cluster.

**Figure 3:**
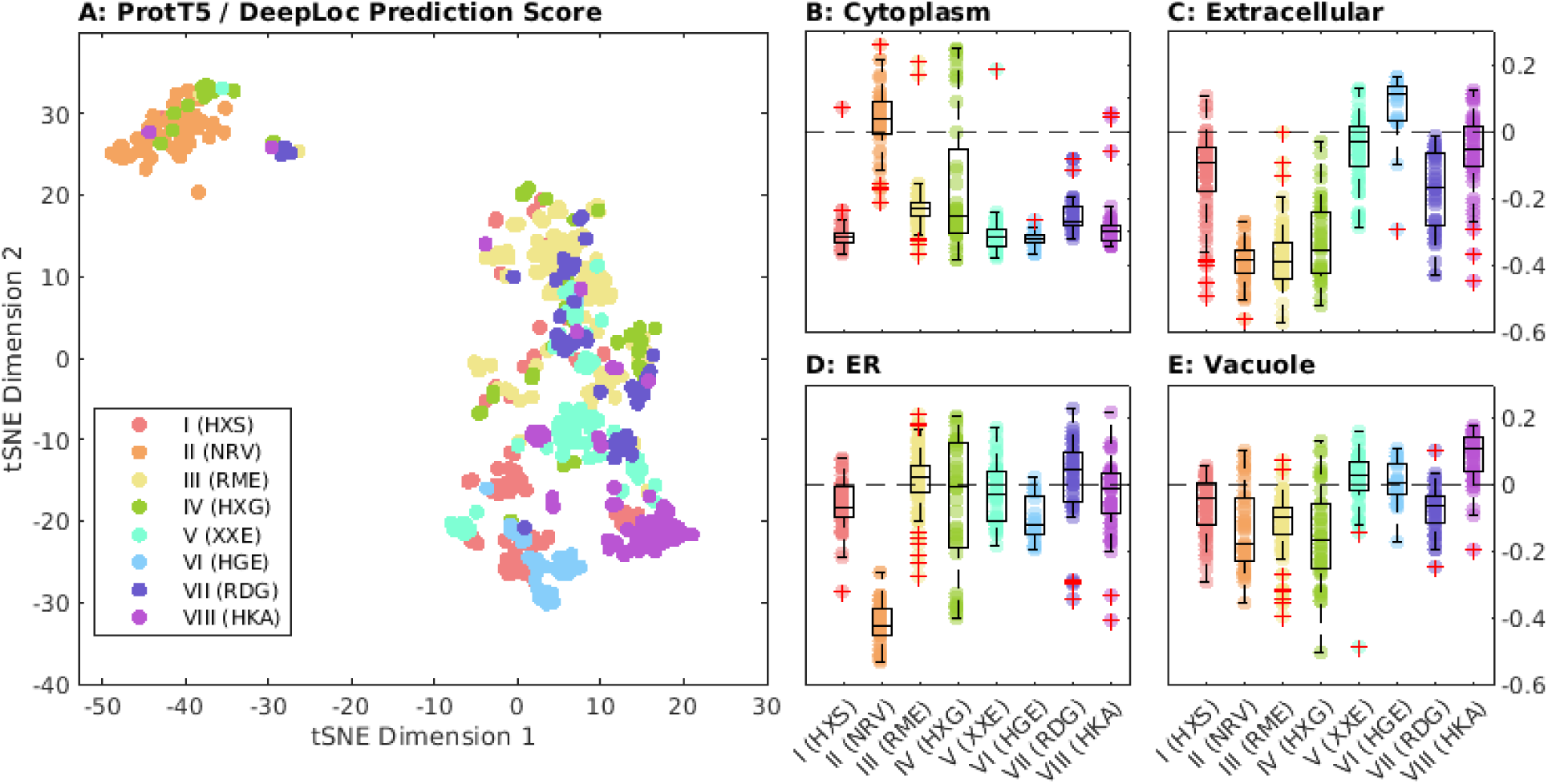
Subcellular localizations of monocotyledonous CKXs predicted by DeepLoc with ProtT5 transformer. **A**: t-Distributed stochastic neighbour embeddings (tSNE) of the DeepLoc prediction scores. Each point represents a single CKX sequence. Each sequence is labelled by its class and the corresponding representative motif of the given class. **B-E**: DeepLoc scores of the individual CKX sequences for the cytoplasm, extracellular space, endoplasmic reticulum (ER), and vacuole. Higher scores indicate higher prediction confidence. Red crosses denote outlying points.

With the non-transformed prediction scores, we also assigned the most likely subcellular localizations to each CKX class. Figure 3B-E shows the class-wise prediction scores for cytoplasm, apoplast, endoplasmic reticulum (ER), and vacuole. Predictions for other compartments were low for all sequences in our dataset. Class II CKXs are predicted to be found mainly in the cytoplasm, classes III and VIII CKXs in the ER, and classes V and VIII CKXs in the vacuole. No predominant localization can be assigned to the class I CKXs, although they are unlikely to be found in cytoplasm. Localizations of class IV CKXs show no clear trend, which is understandable given their lack of clustering in Figure 3A.

### CKX is Active in Xylem Sap and Displays Substrate Specificity Characteristic for Its Root-Derived Counterpart

We measured CKX activity in xylem sap samples from the maize, oat, barley, and wheat. The results are shown in Figure 4. We found the highest CKX activities in maize and oats. We chose the latter for further experiments for two reasons: manipulation with oat plants is less demanding, and we could detect CKX activity in oat xylem sap as early as twelve days after germination. Measuring the CKX activity in the xylem sap at different pH values revealed that the pH optimum of the CKX is 8.5, while the pH of the xylem sap is 6.1.

**Figure 4:**
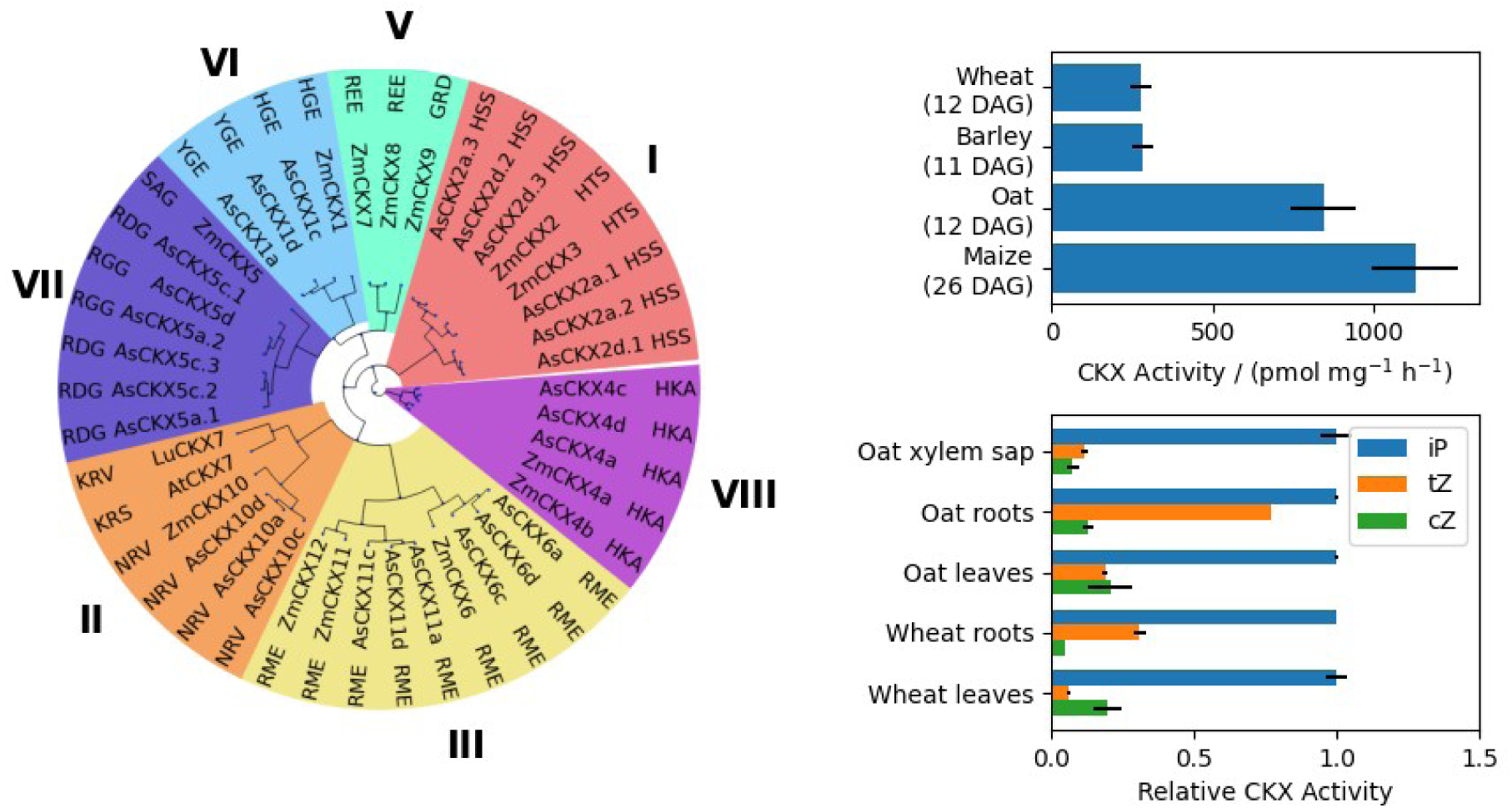
Cytokinin dehydrogenase in the oat (Avena sativa). **Left**: A dendrogram depicting classification of oat and maize CKXs, alongside LuCKX7 and AtCKX7. For each isoform, the three letter-long entrance motif is shown. The classes are annotated by Roman numerals placed around the graph. **Top right**: CKX activity measured in the xylem saps of different plant species. The ages of the plants are given in days after germination (DAG). **Bottom right**: Substrate specificities of CKXs isolated from different parts of oats and maize. iP: isopentenyladenine, tZ: trans-zeatin, cZ: cis-zeatin.

Next, we determined the specificity of the CKX from the oat xylem sap towards iP, tZ, and cZ, i.e. three different CK nucleobases. For comparison, we also performed these measurements with CKXs in samples isolated from the roots and leaves of the oat and wheat. The results are summarized in Figure 4.

In all sample types, iP was the preferred substrate. CKXs obtained from the oat and wheat roots also favoured tZ over cZ. The same trend applied to CKX from the xylem sap, albeit the difference in activities towards the two zeatins was smaller. Conversely, the leaf CKXs preferred cZ over tZ. These findings led us to hypothesize that the CKXs in the oat xylem sap are synthesized in the roots.

Previous substrate specificity studies on maize CKXs have shown that cZ is preferred over tZ by class V ZmCKX8,9 and class II ZmCKX10; conversely, class VI ZmCKX1 favours tZ over its *cis* counterpart (Šmehilová *et al*., 2009; Zalabák *et al*., 2014). Considering the reported extracellular localization of ZmCKX1 (Šmehilová *et al*., 2009; Zalabák *et al*., 2016), we hypothesized that its oat homologues are responsible for CKX activity in xylem sap.

### Oat Genome Harbours CKX Genes with Distinct Expression Patterns in Roots and Leaves

To provide evidence for the putative role of ZmCKX1 oat homologues in CKX activity in the xylem sap, we first needed to identify and characterize oat *CKX* genes overall. In the annotated oat genome, we found 36 putative *CKX* genes (Kamal *et al*., 2022; Tinker *et al*., 2022). Four of them can be expressed in two alternatively splicing variants. Having excluded likely pseudo-genes, we narrowed this list to 27 *AsCKX* isoforms and named them based on their homology with ZmCKXs (see Material and Methods for details). We performed a multi-sequential alignment of the corresponding AsCKX proteins with their maize counterparts. The dendrogram in Figure 4 shows that AsCKXs belong to CKX classes I, II, III, VI, VII, and VIII; ZmCKXs additionally include class V CKXs. We found three oat homologues of ZmCKX1, namely AsCKX1a,c,d; all three belong to class VI and have a glutamate at the VEGAS position.

We have previously hypothesized that the oat homologues of ZmCKX1, i.e. AsCKX1s, are mainly synthesized in roots and secreted into xylem sap. To support this hypothesis, we quantified publicly available *AsCKX* transcript reads in oat leaves, stems, roots, and the pooled samples. In addition, we performed RNA sequencing in oat leaf blades and included the data obtained in this way in our analysis. Relative expressions of *AsCKXs* in all samples are depicted in Figure 5. The plot shows that *AsCKX1s* and *AsCKX4s* are the dominant *CKX* isoforms expressed in roots, followed by three class I *AsCKX2s*. The most expressed forms in the leaves and stems were class II *AsCKX10s*, followed by the remaining *AsCKX2s*, some class III *AsCKX6s*, and *AsCKX4s*. As already implied, the six *AsCKX2s* display two distinct expression patterns. *AsCKX2a.1,a.2,d.1* are slightly expressed in leaves, while *AsCKX2a.3,d.2,d.3* in stems and roots. The latter are expressed slightly more than the former in the pooled samples. The expression profiles shown in Figure 5 support the hypothesis that AsCKX1s are secreted from the roots into the xylem.

**Figure 5:**
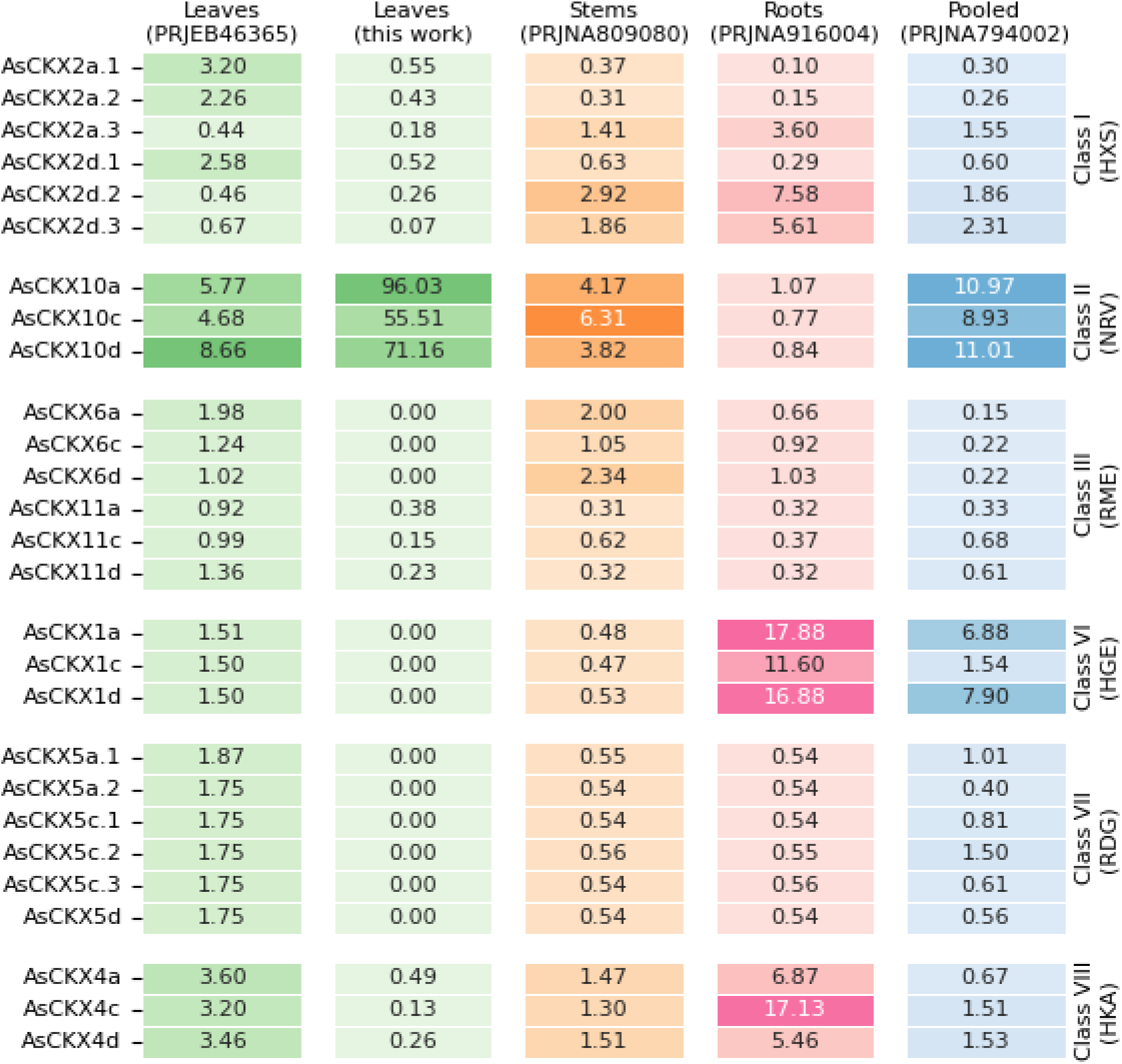
AsCKX expression patterns in oat leaves, stems, roots, and pooled samples. The data represent means of normalized counts obtained via RNA sequencing, and values are given in transcripts per million. The data were obtained either in this work or from publicly available sequence reads; in the latter case, the corresponding project accessions are provided in the column headers. The AsCKX genes are further annotated with the CKX classes and the corresponding characteristic entrance motifs.

### AsCKX Proteins in Oat Xylem Sap are Glycosylated

Another characteristic feature of ZmCKX1, besides its subcellular localization and substrate specificity, is its glycosylation pattern. ZmCKX1 is *N*-glycosylated at six positions with 3-25 hexose-long oligosaccharides. This glycosylation enhances the enzyme activity and thermal stability of ZmCKX1 (Malito *et al*., 2004; Kopečný *et al*., 2008, 2010; Franc *et al*., 2012). We used Concanavalin A Sepharose 4B chromatography to determine the glycosylation of the CKX in oat xylem sap. Most CKX activity in the xylem sap (∼ 87 %) was associated with the glycosylated isoforms (Figure 6).

**Figure 6:**
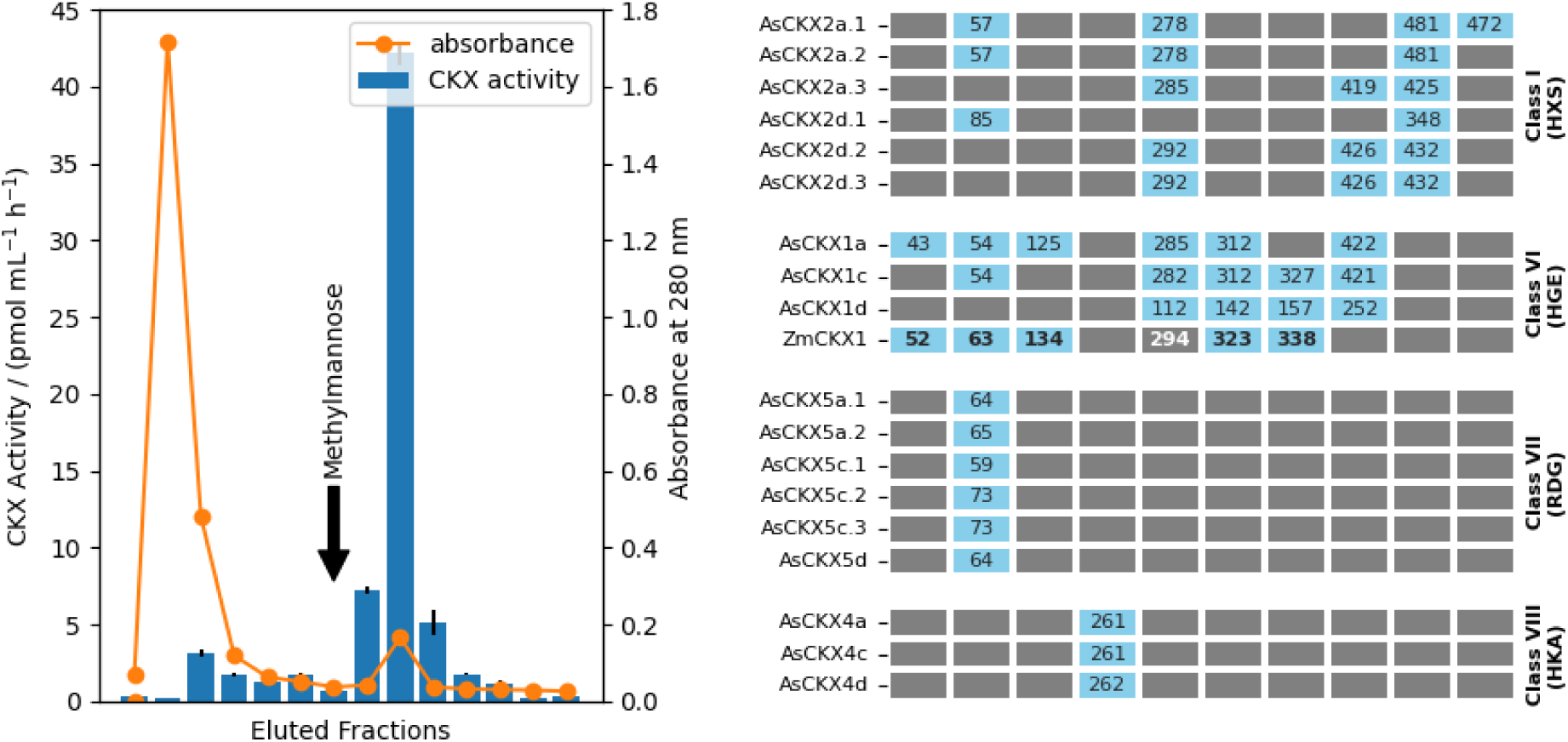
Glycosylation of oat CKXs. **Left**: Absorbance at 280 nm and CKX activity measured in fractions eluted during Concavalin A-Sepharose 4B chromatography. The arrow annotates elution of glycosyated proteins by applying 200 mM methylmannose. **Right**: Prediction of N-glycosylation sites for AsCKXs and ZmCKX1 via NetNGlyc. The light blue cells represent asparagine residues (identified by their numbers given inside the cells) predicted as N-glycosylated with at least 50 % confidence. Residues in each column are aligned among all CKX sequences included in the graphics. For ZmCKX1, the bold numbers represent residues that were experimentally identified as N-glycosylated. CKX classification and the representative entrance motifs are shown on the right side of the graphics. AsCKXs from classes II and III have been excluded, as they are not predicted to be N-glycosylated.

To see whether our hypothesis about the activity of AsCKX1s in the xylem sap is consistent with these findings, we used NetNGlyc software (Gupta and Brunak, 2002) to predict *N*-glycosylation sites in the sequences of AsCKXs and ZmCKX1, for which we could compare the predicted sites with those determined experimentally (Franc *et al*., 2012). The prediction results are summarized in Figure 6.

Class VI AsCKX1s and ZmCKX1 contain 4-6 predicted *N*-glycosylation sites, compared to 2-4 sites for class I, one site for class VII and VIII AsCKXs, and no site for class II and III AsCKXs. The seemingly missing sites in the AsCKX1d sequence might be due to the incomplete annotation of the *AsCKX1d* gene, which is also indicated by the relatively shorter length of the AsCKX1d protein sequence (see Table S3). NetNGlyc predicted five of the six experimentally *N*-glycosylation sites of ZmCKX1. The only exception was N294, although the corresponding residues in some AsCKXs were predicted as *N*-glycosylated with satisfactory confidence.

Two predicted *N*-glycosylation sites are conserved among class II and VI CKXs, and one additional site among CKXs of classes II, VI, and VII. Interestingly, the different expression patterns of AsCKX2s (Figure 5) also reflect the predicted *N*-glycosylation patterns. The one *N*-glycosylation site predicted for class VIII AsCKXs is unique for them.

These predictions indicate a broad glycosylation of AsCKX1s. Together with the data on CKX glycosylation in the xylem, they support the hypothesis that AsCKX1s are mainly responsible for the CKX activity in the xylem sap.

### The CKX Activity in the Xylem Sap Responds to the Nitrate Supply and is Linked to the CK Contents

To further investigate the physiological relevance of CKX in oat xylem sap, we examined changes in CKX activity in the xylem in response to differential nitrate supply, which is known to reflect in CK root-to-shoot flux (Takei *et al*., 2001, 2004; Poitout *et al*., 2018; Sakakibara, 2021). To this end, we grew oat plants at various concentrations of external nitrate (16.0, 62.5, 250.0, and 1000.0 µM) and subjected them to analyses described below.

Firstly, to ensure that the difference in nitrate supply in the nutrient solution corresponds to the nitrate amounts within the oat plants, we measured nitrate concentrations in oat roots, leaves, and xylem sap, as well as the enzyme activity of nitrate reductase (NIA) in roots and leaves. The results (Figure 7F,G) show that both the nitrate concentration and the NIA activity increase with the external nitrate supply.

**Figure 7:**
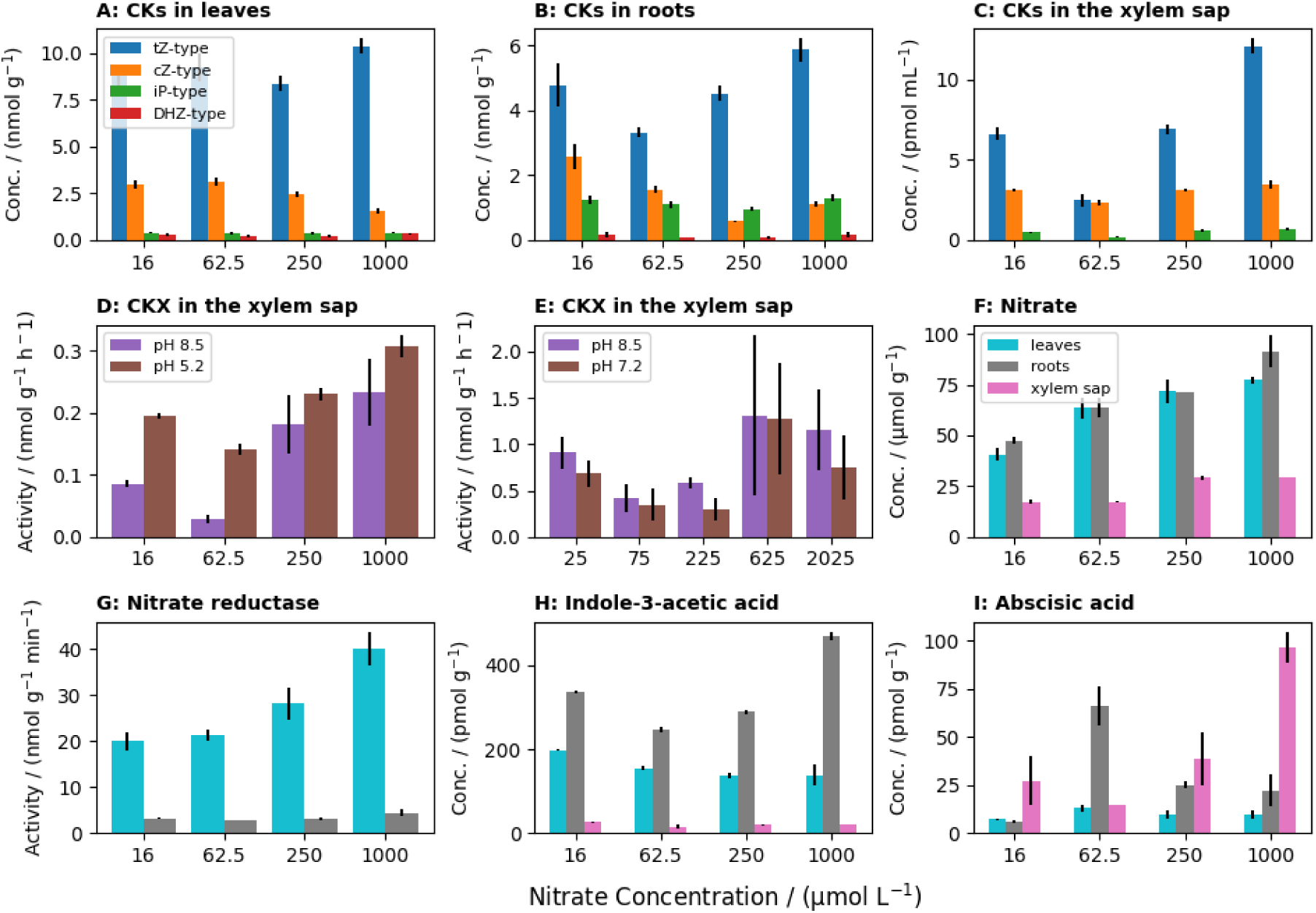
Various responses to different nitrate supply in oats. **A**: CK concentrations in leaves. **B**: CK concentrations in roots. **C**: CK concentrations in xylem sap. **D**: CKX activity in xylem sap at two pH values (nitrate supply ranging from 16 to 1000 µM). **E**: CKX activity in xylem sap at two pH values (nitrate supply ranging from 25 to 2025 µM). **F**: Nitrate concentrations in leaves, roots, and xylem sap. **G**: Nitrate reductase activity in leaves and roots. **H**: Indole-3-acetic acid concentrations in leaves, roots, and xylem sap. **I**: Abscisic acid concentrations in leaves, roots, and xylem sap. tZ: trans-zeatin, cZ: cis-zeatin, iP: isopentenyladenine, DHZ: dihydrozeatin.

Next, we measured concentrations of different CK types (grouped according to the chemistry of their side chains) in the oat xylem sap, roots, and leaves. The results are shown in Figure 7A-C. In each measurement, tZ-type CKs were the most abundant, followed by cZ-type, iP-type, and DHZ-type. The last was detected in the roots and leaves but not the xylem sap.

The concentrations of the tZ-type CKs visibly responded to the external nitrate. However, this response was not monotonous. When the nitrate concentration increased from 16.0 to 62.5 µM, the concentration of tZ-type CK in the xylem sap and roots decreased. With further increase in the nitrate concentration, the concentration of tZ-type CKs grows again, eventually surpassing the concentration measured at 16.0 µM nitrate.

Besides the CKs, we also examined concentrations of two other plant hormones, auxin indole-3-acetic acid (IAA; Figure 7H) and abscisic acid (ABA; Figure 7I). IAA behaved similarly to CKs, with the lowest concentrations at 62.5 µM nitrate. ABA showed a similar trend in the xylem sap but a reverse trend in the roots and leaves (i.e. highest concentrations at 62.5 µM nitrate).

Finally, we determined the CKX activity in the oat xylem sap. Figure 7D shows that when measured at two pH values (8.5 and 5.1), the CKX activity followed the same trend as previsouly observed for CK concentrations in the xylem sap (i.e. an initial drop to a minimum followed by an increment). When we repeated the measurements with a different set of nitrate concentrations (25.0, 75.0, 225.0, 625.0, and 2025 µM), we obtained similar results (Figure 7E). These findings indicate that the CKX activity in oat xylem sap reflects CK concentration and nitrate availability and that CK degradation in the xylem is thus regulated.

## Material and Methods

### Bioinformatical Analysis of CKX Sequences

Known monocot CKX sequences were retrieved from the National Center for Biotechnology Information (NCBI) Identical Protein Groups database (Sayers *et al*., 2022). Oat (*Avena sativa*) CKX sequences were retrieved from the annotated oat cv. Sang reference genome v1.1 (Kamal *et al*., 2022; Tinker *et al*., 2022). We selected genes annotated by gene ontology term GO:0019139 (Ashburner *et al*., 2000; The Gene Ontology Consortium *et al*., 2023) or by human-readable descriptors containing the sub-strings “cytokinin” followed by “dehydrogenase” or “oxidase”. Conserved domains in the selected genes were translated from nucleotide to protein sequences using the ExPASy translate tool (Gasteiger *et al*., 2003).

Oat CKXs were named according to their homology with maize (*Zea mays*). Because of the smaller number of ZmCKXs in comparison with AsCKXs, each AsCKX was also identified with the letter “a”, “c”, or “d”, according to its membership in one of the three *A. sativa* chromosome groups (Tomaszewska, Schwarzacher and Heslop-Harrison, 2022). AsCKXs sharing the closest maize homologue and the chromosome group were further annotated with numbers ranging from one onwards.

All multi-sequence alignments presented in this work were generated using MAFFT software (Katoh and Standley, 2013). Tree graphs were generated from the multi-sequence alignment files using IQ-TREE software (Nguyen *et al*., 2015) and visualized using the Python package ETE (Huerta-Cepas, Serra and Bork, 2016).

### Sequence-Based Predictions

The subcellular localizations of CKX proteins were predicted using DeepLoc (Thumuluri *et al*., 2022) and ProtT5-XL-Uniref50 transformer (Elnaggar *et al*., 2022). For visualization, the prediction scores were transformed from ten-dimensional to two-dimensional space via t-distributed stochastic neighbour embedding (van der Maaten and Hinton, 2008). Glycosylation patterns of CKS proteins were predicted using NetNGlyc (Gupta and Brunak, 2002).

### RNA Sequencing and Quantification

Total RNA was isolated from 50–100 mg of the plant material using FavorPrep Plant Total RNA Purification kit (Favorgen) and treated with rDNAse (Macherey-Nagel). RNA purity, concentration and integrity were assessed with the RNA Nano 6000 Assay Kit using a Bioanalyzer instrument (Agilent Technologies). For RNA-seq analysis, approximately 5 µg of RNA were submitted for the service procedure provided by the Institute of Applied Biotechnologies. The analysis yielded at least 15 million 150 bps read pairs. Publicly available raw transcripts were accessed via the NCBI Sequence Read Archive (Sayers *et al*., 2022) and the European Nucleotide Archive (Leinonen *et al*., 2011). Namely, we used data from projects PRJEB46365 (runs ERR6323384-86), PRJNA916004 (runs SRR22937051-62), PRJNA809080 (runs SRR18094446-57), and PRJNA794002 (runs SRS11553144-65).

Rough reads were quality-filtered using Rcorrector and Trim Galore scripts (Song and Florea, 2015). Transcript abundances were determined using Salmon (Patro *et al*., 2017) with options –-posBias, –-seqBias, –-gcBias, –-numBootstraps 30. The reference index was built from the *A. sativa* v1.1 transcript dataset.

### Crystallization and Structure Refinement of ZmCKX5

The *ZmCKX5* gene (Phytozome ID Zm00001d008862) was cloned into a pTYB12 vector and expressed in T7 Express *Escherichia coli* cells (New England Biolabs) at 18 °C overnight. Protein was purified by chitin-based affinity chromatography upon elution with 50 mm DTT to cleave the intein tag and further by gel permeation chromatography on a Superdex S200 column using the NGC liquid chromatography system (Bio-Rad). ZmCKX5 (23 mg mL-1) in 50 mM Tris-HCl, pH 8.0, was crystallized in the condition found using the NeXtal PEGII Suite (Qiagen) containing 0.2 M Li_2_SO4, 0.1 M Tris-HCl pH 8.5, 30 % PEG 3000. Crystals were cryoprotected with 15 % glycerol and flash-frozen in liquid nitrogen. Diffraction data were collected at 100 K on the Proxima 2 beamline at the SOLEIL synchrotron (Saint-Aubin, France) at 1.65 Å resolution. Intensities were integrated using the XDS program (Kabsch, 2010). Data quality was assessed using the correlation coefficient *CC*1/2 (Karplus and Diederichs, 2012). The crystal structure of ZmCKX5 was determined by performing molecular replacement with Phaser (Storoni, McCoy and Read, 2004) using a monomer of ZmCKX1 (PDB ID 2QKN) as a search model (Kopečný *et al*., 2010). The model was refined with BUSTER-TNT (Bricogne *et al*., 2017), and electron density maps were evaluated using COOT (Emsley and Cowtan, 2004). Structure quality was validated using MolProbity (Chen *et al*., 2010). Molecular graphics were generated using PyMOL software (Schrödinger, LLC, 2015).

### Plant Material and Growth Conditions

For the determination of nitrate and CK concentrations and CKX activity, oat plants were grown hydroponically for 12 days in a growth room with 21 °C/15 °C day/night air temperature and 16 h photoperiod (photon flux of 400 µmol m^-2^ s^-1^). Continuously aerated nutrient solution contained 158 µM Ca(NO_3_)_2_, 70.8 µM KNO_3_, 52.5 µM KH_2_PO_4_, 41.3 µM MgSO_4_, 47.5 µM KCl, 2.5 µM H_3_BO_3_, 2 µM Fe-EDTA, 0.2 µM ZnSO_4_, 0.2 µM MnSO_4_, 0.05 µM CuSO_4_, and 0.01 µM (NH_4_)_6_Mo_7_O_24_. For each treatment, 200 L of nutrient solutions were used, corresponding to 2 L per plant. The nutrient solution was changed weekly, and the nitrate concentration was checked and adjusted every other day. Two days before the sampling, the plants were twice transferred to fresh nutrient media containing 16.0, 62.5, 250.0 or 1000.0 µM nitrate and kept there for a day. Missing Ca^2+^ and K^+^ were supplied in the form of CaCl_2_ and KCl.

For other experiments, plants were soaked for 14 h in aerated distilled water and then sown on perlite saturated with double-concentrated Knop’s nutrient solution. Plants were grown in a controlled climate growth chamber (Sanyo MLE-350H) at 20 °C/18 °C day/night temperature, 80 % air humidity, and 16 h photoperiod with a photon flux density of 300 µmol m^-2^ s^-1^. Unless stated otherwise, plants were grown for 12 days or until they had two fully developed leaves and one emerging.

### Sampling of Xylem Sap, Roots and Shoots

For xylem sap collection, shoots were excised with a razor blade approximately 0.5 cm above the shoot-to-root transition. The xylem sap released during the first 15 min was discarded. During the following two hours, the sap drops were collected, immediately cooled and kept on ice in closed tubes. Collected sap samples were frozen in liquid nitrogen and stored at –80 °C. Shoots and roots were frozen in liquid nitrogen and stored under the same conditions.

### Determination of Nitrate Content

Leaf and root samples (1 g of fresh weight) were homogenized in liquid nitrogen, extracted with deionized water for 30 min at 90 °C, and filtered. Nitrogen concentrations in plant extracts, nutrient solutions, and xylem sap were determined spectrophotometrically using Skalar San plus analyser (Breda, the Netherlands). The samples were passed through a column of granulated copper-cadmium to reduce nitrate to nitrite. The nitrite was determined spectrophotometrically by measuring the conversion of sulfanilamide and *N*-(1-naphthyl)ethylenediamine dihydrochloride to an azo dye at 540 nm.

### Measurement of NIA Activity

Leaf and root samples (1 g of fresh weight) were homogenized in liquid nitrogen and extracted in 5 mL of 50 mM Tris-HCl buffer, pH 8.0, containing 3 % bovine serum albumin at 4 °C for 30 min. Insoluble material was removed via centrifugation (1500 × g, 30 min). The NIA activity was determined by measuring the conversion of nitrate to nitrite according to (Gaudinová, 1983).

### Cytokinin Analysis

CK-containing fractions were isolated from plant samples via dual-mode solid phase extraction (Dobrev and Kamínek, 2002). CK detection and quantification were carried out using LC/MS/MS system consisting of HTS-Pal auto-sampler with a cooled sample stack (CTC Analytics, Zwingen, Switzerland), Rheos 2200 quaternary HPLC pump (Flux Instruments, Basel, Switzerland), Delta Chrom CTC 100 Column oven (Watrex, Prague, Czechia), and TSQ Quantum Ultra AM triple-quad high-resolution mass spectrometer (Thermo Electron, San Jose, USA) equipped with an electrospray interface. The mass spectrometer was operated in the positive single-reaction monitoring mode.

### Measurement of CKX Activity

The CKX activity was measured by an *in vitro* radioisotope assay based on the conversion of [2-^3^H]iP (prepared by the Isotope Laboratory of the Institute of Experimental Botany AS CR, Prague, Czechia) to [^3^H]adenine (Motyka *et al*., 2003). The xylem exudate (20 μL per assay) was used without previous purification. Protein concentration in the xylem sap was determined according to Bradford (1976) using bovine serum albumin as a standard. The substrate and the product were separated as described by Gaudinová et al. (2005). To determine CKX substrate specificity, [2-^3^H]iP was replaced with [2-^3^H]tZ or [2-^3^H]cZ in the standard assay mixture (Gajdošová *et al*., 2011). All radio-labelled substrates were used at a concentration of 2 µM and a molar activity of 7.4 Bq mol^-1^.

### Determination of CKX Glycosylation

The glycosylation state of CKX in oat xylem sap was determined by Concanavalin A-Sepharose 4B chromatography (Motyka and Kamínek, 1994; Motyka *et al*., 1996). In the collected fractions, CKX activity was determined as described above.

## Discussion

In this work, we address the multifaceted diversity of CKX as a conserved trait enabling selective and fine-tuned regulation of CK homeostasis. Focusing on monocots, we divided 492 CKXs into eight classes based on their sequential similarity. This CKX classification correlates with the variance of the VEGAS residue-containing “entrance triplet”, i.e. the three consecutive variable residues of the FL**X**RV**XXX**E motif (with VEGAS being the third of them).

This correlation suggests that sequentially closely related CKX isoforms are similar in substrate specificities and enzyme activities. Given that the species included in our analysis possess several CKX isoforms belonging to different classes (Figure 1), the individual isoforms might play specific roles in the regulation of the CK metabolism rather than being simply redundant (although redundancy among isoforms belonging to the same class remains possible). This hypothesis goes along with the differential affinities of maize CKXs towards various CK substrates. ZmCKX2,3 (class I) and ZmCKX4a,b (class VIII) have shown preference towards CK nucleotides and N9-glucosides, while ZmCKX12 (class III), ZmCKX8,9 (class V), and ZmCKX1 (class VI) preferred CK nucleobases, and ZmCKX10 (class II) and ZmCKX5 (class VII) cleaved some N9-derived CKs similarly to the nucleobases. (Šmehilová *et al*., 2009; Zalabák *et al*., 2014; Kopečný *et al*., 2016).

The CKX substrate specificity about the N9 substitution has been attributed to the varying character of the VEGAS residue and supported by experimentally solved structures of CKXs from different classes (Malito *et al*., 2004; Bae *et al*., 2008; Kopečný *et al*., 2016; Wan *et al*., 2019; Nisler *et al*., 2021). In this work, we have also solved the structure of ZmCKX5, thus covering six of the eight CKX classes. The superimpositions depicted in Figure 2 confirm the unique capacity of a glutamate residue to interact with a nucleobase substrate, but they also reveal the variability of a non-conserved flexible segment close to the entrance tunnel. This segment might further regulate the substrate specificity and enzyme activity through its composition and conformation, with the latter capable of restricting the access of the substrate to the active site.

CKXs also differ in their specificities towards CKs with varying characters of the N6 substituent (i.e. the side chain). For instance, ZmCKX8,9 strongly preferred *cis*-(cZ) and *trans*-zeatin (tZ), both CKs with a hydroxylated side chain, over non-hydroxylated isopentenyladenine (iP). Conversely, ZmCKX1 preferred iP and tZ over cZ, and ZmCKX10 cleaved all three substrates at similar rates (Šmehilová *et al*., 2009; Zalabák *et al*., 2014, 2016; Kopečný *et al*., 2016). All these proteins have glutamate at the VEGAS position but belong to different classes, showing that the sequence-based substrate specificity of CKX goes beyond the VEGAS residue. The composition of the entrance triplet likely co-relates with other sequential (and consequently structural) features responsible for additional fine-tuning of the protein’s substrate specificity. The ability of CKXs to discriminate CKs with different N6 and N9 substitutions could also contribute to the varying effects of *OsCKX* mutations on endogenous concentrations of tZR, iP, and iPR in rice (Zheng *et al*., 2023). Our results suggest that the diversity of CKX isoforms and their substrate specificities is conserved among the monocot species, if not the plant kingdom as a whole, considering that similar trends were observed in *Arabidopsis* (Frébortová *et al*., 2007; Galuszka *et al*., 2007; Kowalska *et al*., 2010).

We also saw a correlation between the CKX classification and the predicted subcellular localizations, further backing the idea of the CKX family consisting of similar yet functionally specialized members. Combining different substrate specificities and subcellular localizations allows us to discuss the role of CKX in the context of the diverse and precisely regulated CK distribution at the cellular level (Nedvěd *et al*., 2021). In this work, we predict that most of the analysed CKXs localize to cytoplasm, ER, vacuole or are secreted. Most cytoplasmic isoforms belong to class II and contain a valine at their VEGAS position – a small and non-polar residue allowing the binding of bulky substrates.

CKXs predicted to localize to the ER mostly belong to classes III and VII, partly also to classes IV and V. A portion of the ER-localized CKXs possesses a glutamate residue at the VEGAS position and might thus efficiently cleave CK nucleobases. They could directly interfere with CK signalling involving ER-localized CK receptors (Wulfetange *et al*., 2011). CKXs predicted to be localized to the vacuole are represented by classes V and VIII, which typically contain a glutamate or alanine residue at their respective VEGAS positions. The vacuole was reported as the final compartment of most ZmCKXs expressed in an *Arabidopsis* cell culture (Zalabák *et al*., 2016). Considering the existence of vacuole-stored CKs (Jiskrová *et al*., 2016), plants might use the CKX-catalysed cleavage to recycle adenine species. Vacuolar CKXs might thus contribute to the export of adenosine from the vacuole to the cytoplasm, which has been described as crucial for growth and pollen germination (Bernard *et al*., 2011).

Class VI CKXs are likely secreted to the apoplast, which makes them good candidates for proteins responsible for the CKX activity in the xylem addressed before (Hoyerová *et al*., 2007). They are also likely to regulate the amount of extracellular CK nucleobases available for plasma membrane-residing CK receptors (Antoniadi *et al*., 2020). Additionally, the class VI representative in maize, ZmCKX1, has repeatedly shown exceptionally high enzyme activity in comparison with the other ZmCKX isoforms (Šmehilová *et al*., 2009; Zalabák *et al*., 2014, 2016). Such highly active and nucleobase-specific CKX might contribute to the lower abundance of CK nucleobases in the xylem sap in comparison to CK nucleosides (Beveridge *et al*., 1997; Takei *et al*., 2001; Sakakibara, 2021).

Subsequent analyses and experiments further supported the hypothesis that CKX activity found in the oat xylem sap (Hoyerová *et al*., 2007) is to be attributed to the oat homologues of ZmCKX1, namely AsCKX1a,c,d. The corresponding *AsCKX1a,c,d* genes are mainly expressed in roots (Figure 5), as we expect in the case of proteins that we consider excreted to the xylem and transported via the transpiration stream to the shoot. CKXs isolated from the oat roots also show a similar pattern of substrate specificity towards iP, tZ, and cZ as their xylem sap counterparts (Figure 4). Furthermore, our measurements revealed that the active xylem sap CKXs are glycosylated proteins, which is consistent with the previously reported glycosylation of ZmCKX1 and its enhancing effect on ZmCKX1 enzyme activity (Franc *et al*., 2012).

As summarized in the Introduction, CKs in the xylem act as messengers carrying information about nitrate availability in the environment (Kiba *et al*., 2011; Ruffel *et al*., 2016; Poitout *et al*., 2018; Roy, 2018; Sakakibara, 2021). It follows that the xylem-located CKX might respond to the nitrate supply as well. To confirm this assumption, we examined the changes in CK concentrations and CKX activity in oat plants exposed to different concentrations of nitrate. In agreement with previous findings (Takei *et al*., 2001; Poitout *et al*., 2018), we observed a significant response to nitrate among tZ-type CKs, which were the most abundant in our samples. The concentrations of these CKs varied in the oat roots and xylem but not in the leaves, suggesting that the leaves possess mechanisms to compensate for the increased CK influx from the roots. In our case, the root-borne CKs might be degraded by AsCKX10s, whose genes are highly expressed in leaves. AsCKX10s belong to class II and therefore likely recognize *trans*-zeatin riboside (tZR), the major xylem-located CK (Takei *et al*., 2001; Osugi *et al*., 2017; Sakakibara, 2021), as a substrate. The ability to compensate for the root-borne CKs in leaves would allow local or systemic shoot-to-root CK signalling to occur without being necessarily influenced by the transpiration flux.

We also show that the nitrate-dependent changes in CK concentrations in the roots and xylem sap correlate with the changes in the activity of xylem-located CKXs, suggesting that CKX in the xylem sap responds to concentrations of transported CKs (and by extension to nitrate supply). This response was apparent when tested at different pH values and nitrate concentration ranges (Figure 7D,E). As described above, we have attributed the xylem-located CKX activity to AsCKX1s, which can cleave biologically active CK nucleobases. Their physiological role might comprise a negative regulation of the CK nucleobases present, thus preventing their leakage along the way. An increase in the xylem-located CKX activity might also contribute to a previously observed nitrogen-induced shift in the tZR/tZ ratio in the xylem (Osugi *et al*., 2017; Sakakibara, 2021).

It is apparent from Figure 7B-E that the responses of root and xylem-located CKs and xylem-located CKX were not monotonous but rather passed through a minimum between the lowest and the highest nitrate concentrations tested. One possible explanation of these trends involves a variable efficiency of nitrate perception depending on the nitrate supply. This property has been attributed to NRT1 / PTR FAMILY 6.3 (NPF6.3) protein, which functions as both a nitrate transporter and receptor (Liu, Huang and Tsay, 1999; Ho *et al*., 2009; Wang *et al*., 2018). When the nitrate supply increases from low to high, NPF6.3 switches from a high-to a low-affinity state. The molecular mechanism of the switch involves the dephosphorylation of a threonine residue (T101) and the formation of an NPF6.3 dimer. When the nitrate availability diminishes, T101 gets phosphorylated, and the NPF6.3 dimer dissociates (Parker and Newstead, 2014; Sun *et al*., 2014; Tsay, 2014; Sun and Zheng, 2015). This mechanism of affinity reversal allows us to hypothesize that with growing nitrate supply, there is a point at which a portion of the NPF6.3 population has switched from the high-to the low-affinity state, and the said growth in nitrate supply is not sufficient to compensate for this change. Such a point would explain the observed minima in CK and CKX responses to the nitrate.

Besides CKs and CKX, we also saw that the differences in nitrate supply affected the concentrations of IAA (an auxin) and ABA in the roots and xylem sap. IAA followed the same trend as CKs (Figure 7H). ABA showed the same trend in the xylem sap but a reversed one in the roots (Figure 7I). The latter suggests an antagonistic relation between ABA and CKs, which might be relevant to the previously reported inhibitory effect of ABA on nitrate uptake (Harris and Ondzighi-Assoume, 2017; Su *et al*., 2021).

To sum up, we report that CKX is active in the xylem sap and regulated in response to the nitrate availability. These findings allow us to incorporate CK degradation into contemporary models of CK long-distance transport and root-to-shoot signalling (Nedvěd *et al*., 2021; Sakakibara, 2021; Hu and Shani, 2023). CK cleavage in the apoplast appears to be a possible manner of suppressing the CK signalling, alongside CK transport into cells. Given our assumption that the xylem-located CKXs preferentially cleave CK nucleobases as their substrates, the exact fate of CK ribosides, the main form of the xylem-located CKs, remains yet to be uncovered.

## Acknowledgements

The authors wish to thank to Stanislav Vosolsobě for his advice on the bioinformatical analyses, Eva Kobzová for her assistance in plant cultivation, and Selgen^®^ company for providing seeds necessary for our experimental work.

## Notes

### Competing Interest Statement

The authors have declared no competing interest.

