## Supplementary material for "Cytokinin Dehydrogenase in Xylem Sap Reveals A Direct Link Between Cytokinin Metabolism and Long-Distance Transport": 2023_Nedved_Motyka_CKX_SUPPLEMENT.docx

### Supplementary Information

#### Supplementary Tables

*Table S1: CKX protein sequences used in the bioinformatical analysis. The “ID” coumn consists of the Sang v1.1 gene names for Avena sativa and the NCBI Protein accessions for other species.*

*Table S2: Refinement statistics for the crystallization of ZmCKX5.*

*Table S3: Naming of the AsCKX isoforms. The corresponding CKX classes, lengths of mRNAs and proteins are given as well.*

#### Other Files

A tree graph in the Newick format used to construct the dendrogram in the Figure 1 is provided in the “TREEFILE.txt” file.
